# Detecting neuroplastic effects induced by ketamine in healthy human subjects: a multimodal approach

**DOI:** 10.1101/2025.05.01.651686

**Authors:** Claudio Agnorelli, Joseph Peill, Gabriela Sawicka, Danielle Kurtin, Ekaterina Shatalina, Kirran Ahmad, Matthew B Wall, Catarina Rua, Kate Godfrey, Natalie Erl, Graham Searle, Katie Zhou, Brandon Weiss, Andrea Fagiolini, Robin Carhart-Harris, Paul M. Matthews, Eugenii A. Rabiner, David Nutt, David Erritzoe

## Abstract

We investigated ketamine’s neuroplastic effects in healthy human subjects using integrated Positron Emission Tomography (PET)/Magnetic Resonance Imaging (MRI) measures before and 1-8 days after a single psychedelic dose of ketamine (1 mg/kg, intravenous). Eleven participants underwent two PET/MRI scans with [^11^C]-UCBJ (synaptic density/plasticity), ^1^H-MRS (Glutamate and GABA), and resting-state fMRI (intrinsic brain activity, functional connectivity, graph-theoretic metrics), before and after ketamine. While group-level analyses showed only trend-level increases in PET synaptic markers, we observed significantly elevated Anterior Cingulate Cortex (ACC) glutamate levels post-ketamine. Functional connectivity analyses revealed decreased within-network integrity, particularly in high-order networks like the default mode network (DMN), alongside increased low-to-high-order network integration. Our multimodal analysis showed that increased [^11^C]-UCBJ volume distribution (VT), a putative index of synaptic plasticity, correlated with reduced intrinsic activity in DMN regions and decreased influence of the posterior cingulate cortex (PCC) in global network dynamics. By linking molecular and network-level changes, our results point to the PCC as a central hub where ketamine may reshape brain hierarchies in the long term, providing new directions for understanding its therapeutic mechanisms and developing targeted treatments.

## 1. Introduction

Ketamine is a versatile compound with a tightly dose-dependent range of psychoactive effects. With increasing dosage, ketamine leads to a progression from disinhibition and alcohol-like effects to psychedelic experiences—termed dissociation in clinical contexts—and, at the highest doses, to an anaesthetic state [^1^]. In recent years, clinical trials have revealed ketamine to have a rapid antidepressant effect that occurs within a few hours and lasts for up to a week after a single dose, with repeated exposures extending the duration significantly [^2^]. These sustained clinical effects observed long after the drug has been metabolized prompt critical questions regarding their underlying mechanisms.

A substantial body of preclinical research has demonstrated that the prolonged mood-enhancing effects of ketamine, as well as other antidepressants like SSRIs and classic psychedelics, are associated with enduring neuroplastic changes [^3^]. Cortical neurons cultured with ketamine exhibit increases in dendritic complexity and spine density, essential components for the genesis of new synaptic connections [^4^]. *In vivo*, studies of rodents have shown that a single psychedelic dose of ketamine promotes neurogenesis in the adult hippocampus as well as an increase in dendritic spines and synaptic density within the pre-frontal cortex (PFC). These enhancements often correlate with improvements in behavioural surrogate markers of depressive symptoms [^5,6^]. Furthermore, ketamine reliably reduces the thresholds for stimulation-induced functional modifications, such as short-term and long-term plasticity, in cortical neurons within hours to days post-administration depending on the dose and stimulation paradigm. These modifications are linked to the re-opening of critical developmental windows, manifesting as shortened functional recovery period following monocular deprivation [^7^] or the reactivation of a sensitive period for social reward learning [^8^].

Deficits in neuroplasticity within mood-regulating brain regions have been recognized as core features of various neuropsychiatric disorders, including depression [^9^]. Recent advances in positron emission tomography (PET) techniques have enabled the *in vivo* quantification of molecular markers of synaptic plasticity in the human brain through the validation of the radioligand [^11^C]-UCBJ, which binds to synaptic vesicle protein 2A (SV2A), a protein widely present in presynaptic vesicles [^10^]. Foundational studies using this tracer have reported significant reductions in cortical SV2A concentrations in individuals with neuropsychiatric/neurodegenerative conditions, including depression [^11–13^].

However, despite considerable animal research indicating that ketamine’s rapid and sustained antidepressant effects stem from increased structural and functional neuroplasticity, confirming these findings in the living human brain presents significant challenges. In 2022, Holmes et al. investigated the changes in synaptic SV2A density induced by ketamine (0.5 mg/kg) using [^11^C]-UCBJ in depressed patients, individuals with trauma, and healthy subjects, but found no significant effects at 24 hours post-administration of ketamine. Notably, a *post hoc* exploratory analysis revealed that ketamine did significantly increase SV2A density in the group of depressed patients with the lowest baseline SV2A levels [^14^].

While not directly related to the classical concept of synaptic plasticity as studied in preclinical research, measures of functional whole-brain dynamics can provide valuable insights into the intrinsic flexibility of neuronal network information processing [^15^]. Evidence from functional magnetic resonance imaging (fMRI), indicates that ketamine induces profound acute and sub-acute alterations in intrinsic brain activity and connectivity both within and between neural networks, resulting in lasting changes in whole-brain dynamics that are similar to those produced by classic psychedelics (for reviews see [^16,17^]). Acutely, ketamine was observed to reduce the brain-wide fractional amplitude of low frequency fluctuations (ALFF), a raw index of intrinsic regional neural activity extracted from blood-oxygen-level-dependent (BOLD) signal [^18^]. However, another study observed increases in fractional ALFF in the posterior cingulate cortex (PCC) at 1 hour post-ketamine in healthy subjects, but these changes were not significant at 1 day post-treatment [^19^]. Many studies have demonstrated that ketamine acutely reduces resting-state functional connectivity (FC) within the default mode network (DMN) [^20,21^] (although not universally reported [^19^]), which is involved in ‘narrative’ self-experience and is hyperactive in depression [^22,23^]. This effect is still present at 1 day after exposure to a single dose of ketamine [^19,24,25^]. Also, ketamine reduced the anti-correlation between the DMN and other networks acutely [^20^] and 1 day after the infusion [^25^]. In depressed individuals, altered connectivity in the DMN and other high-order networks was found to correlate with improvements in depression symptomatology [^26–29^].

Collectively, existing literature suggests that ketamine may induce neuroplastic changes linked to mood improvements in humans across multiple levels. However, a comprehensive characterization of ketamine’s neuroplastic effects across different scales, from synaptic alterations to global brain dynamics, and the interactions between these levels remains lacking. Employing innovative multimodal neuroimaging paradigms, recent investigations have revealed intriguing relationships between molecular, cellular, and whole-brain functional dynamics, with functional relevance for neuropsychiatric conditions [^30^]. Using magnetic resonance spectroscopy (MRS) imaging integrated with PET, Onwordi and colleagues (2021) observed a positive correlation between baseline concentrations of glutamate and SV2A, measured via [^11^C]-UCBJ, in the ACC and hippocampus of healthy subjects[^31^]. In another investigation, greater synaptic density in both the PFC and striatum was associated with greater fractional ALFF of the DMN in healthy subjects [^32^]. Examining the impact of synaptic loss on brain network function in frontotemporal lobar degeneration syndrome, it was reported that lower [^11^C]-UCBJ binding correlated with diminished functional connectivity, which was further linked to clinical severity [^33^]. Yet, despite these advances in multimodal neuroimaging techniques, there is currently very limited evidence on how pharmacological interventions, particularly ketamine, influence the relationship between molecular and whole-brain functional dynamics.

In the present study, we employed integrated PET/MRI using [^11^C]-UCBJ, measured from 1 to 8 days after a single psychedelic dose of ketamine (1 mg/kg) in 11 healthy subjects, to assess its effects across different levels of brain function. We hypothesized that ketamine would induce increased density of the SV2A marker of synaptic plasticity, measured via [11C]-UCBJ volume distribution (VT), in mood-regulating regions, with effects emerging up to a week following administration. The relationship between synaptic plasticity and brain function was explored through multimodal neuroimaging using a data-driven approach, including measures of glutamatergic neurotransmission (^1^H-MRS), regional brain activity (ALFF), and whole-brain functional connectivity. Ketamine was expected to decrease regional intrinsic neural activity and within-network integrity while enhancing between-network connectivity, with these changes correlating with increases in synaptic plasticity.

## 1. Materials and methods

### 1.1. Participant recruitment and screening

A total of 11 healthy male participants (mean age 32 ± 10 years) were screened and included in the study. Participant’s details are listed in Table 1. The inclusion criteria were as follows: participants needed to be between 20 and 60 years of age, exhibit no physical or psychiatric medical conditions, and have no history or current incidence of substance abuse. Consumption of ketamine or classic psychedelics in the 6 months preceding the beginning of the study was a ground for exclusion to avoid carry-over neuroplastic effects from previous substance exposure [^3^]. Of the total sample, none of the participants were naïve to classic psychedelics, while n = 7 participants were ketamine naïve. Additionally, participants were required to abstain from alcohol and illicit substance intake for at least 1 week before and throughout the study period. Participants were screened for contraindications to MRI or research PET scanning. Ethical approval was granted by the Brent Research Ethics Committee, London, UK (Supplementary Data).

**Table 1:**
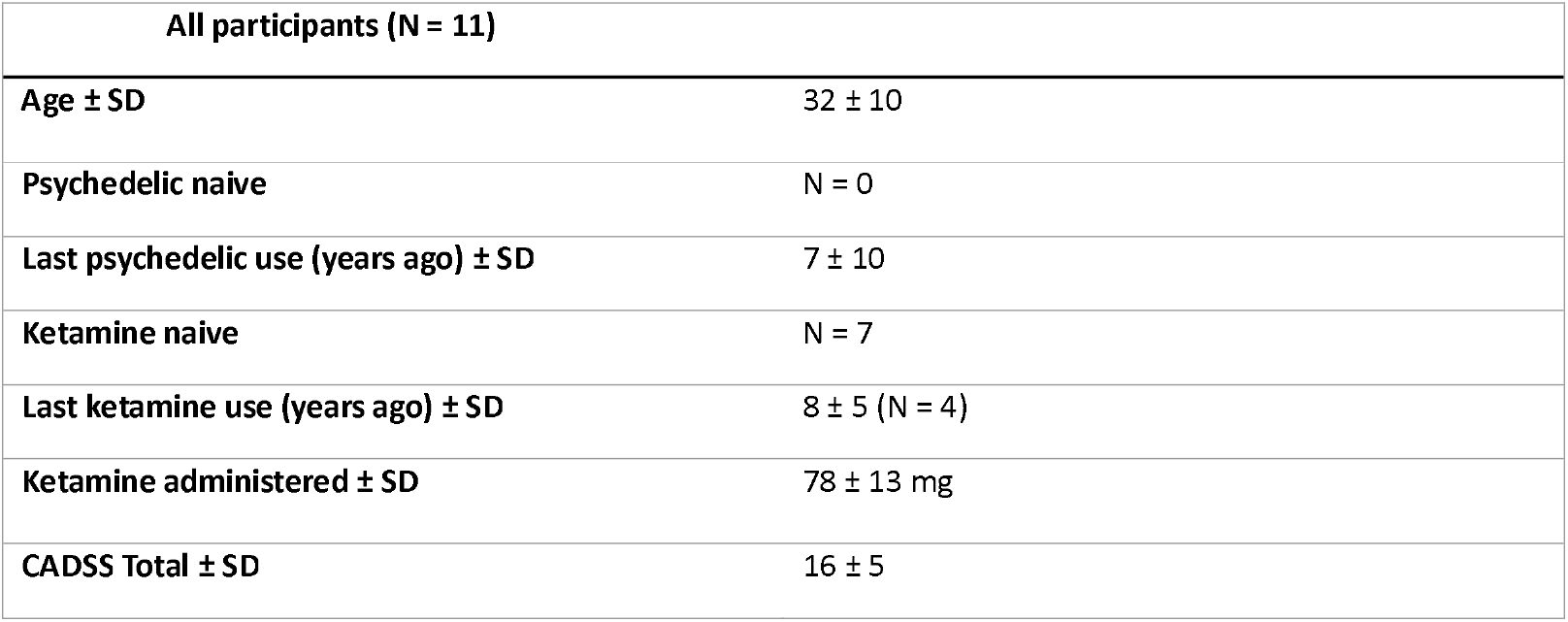
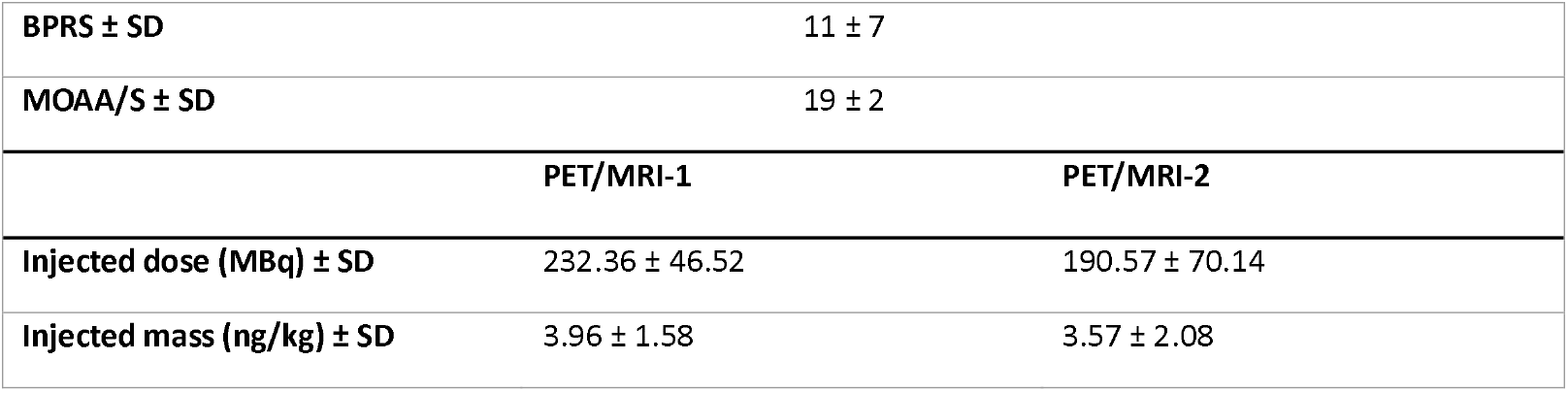
Demographic, drug, and radiotracer characteristics for all participants. SD = standard deviation.

| All participants (N = 11) |  |
| --- | --- |
| Age ± SD | 32 ± 10 |
| Psychedelic naïve | N = 0 |
| Last psychedelic use (years ago) ± SD | 7 ± 10 |
| Ketamine naïve | N = 7 |
| Last ketamine use (years ago) ± SD | 8 ± 5 (N = 4) |
| Ketamine administered ± SD | 78 ± 13 mg |
| CADSS Total ± SD | 16 ± 5 |

|  |  |  |
| --- | --- | --- |
| **BPRS ± SD** | 11 ± 7 | |
| **MOAA/S ± SD** | 19 ± 2 | |
|  | **PET/MRI-1** | **PET/MRI-2** |
| **Injected dose (MBq) ± SD** | 232.36 ± 46.52 | 190.57 ± 70.14 |
| **Injected mass (ng/kg) ± SD** | 3.96 ± 1.58 | 3.57 ± 2.08 |

### 1.2. Ketamine administration

Racemic ketamine was obtained from the St Charles Centre for Health & Wellbeing Pharmacy and administered intravenously (i.v.) by constant infusion over 40 minutes at a dose of 1 mg/kg. This dosage is subanaesthetic and produces a psychedelic-like experience often described as dissociative, and has confirmed antidepressant effects [^34^]. The amount of ketamine administered was comparable across participants (78 ± 13 mg), and all participant experienced the dissociative effects of the drug (Table 1). Vital signs (blood pressure and heart rate) were obtained before, during, and after ketamine infusion. The psychological safety of participants during the acute effects of ketamine was assessed through the Brief Psychiatric Rating Scale (BPRS) [^35^], the Modified Observer’s Assessment of Alertness/Sedation (MOAA/S) scale [^36^], and the 6-item version of the Clinician-Administered Dissociative States Scale (CADSS) [^37^], all administered by the study physician following drug administration.

### 1.3. Study design

Each participant underwent 2 PET/MRI scans, where [^11^C]-UCBJ was administered intravenously for the quantification of brain SV2A (Figure 1A). Subsequently, ^1^H-MRS was acquired to quantify glutamate and GABA concentrations within the ACC. Then, resting-state fMRI data were obtained to assess whole-brain intrinsic brain activity and connectivity. Participants had their first scan (i.e., PET/MRI-1) approximately two weeks before the ketamine infusion. Of the 11 participants, a subset of 4 participants underwent their post-ketamine scan (i.e., PET/MRI-2) 1 day after ketamine infusion. One participant had the scan 2 days after ketamine infusion due to tracer production failure 1-day post-ketamine administration. Participants who had their second scan either at 1 or 2 days after ketamine are referred to as the Day 1-2 group (n= 5). A subset of 4 participants had their Scan 2 at 7 days after ketamine infusion. Two participants had their Scan 2 at 8 days after ketamine infusion due to tracer production failure at 7 days post-ketamine administration. Those participants are referred to as the Day 7-8 group (n = 6).

**Figure 1.**
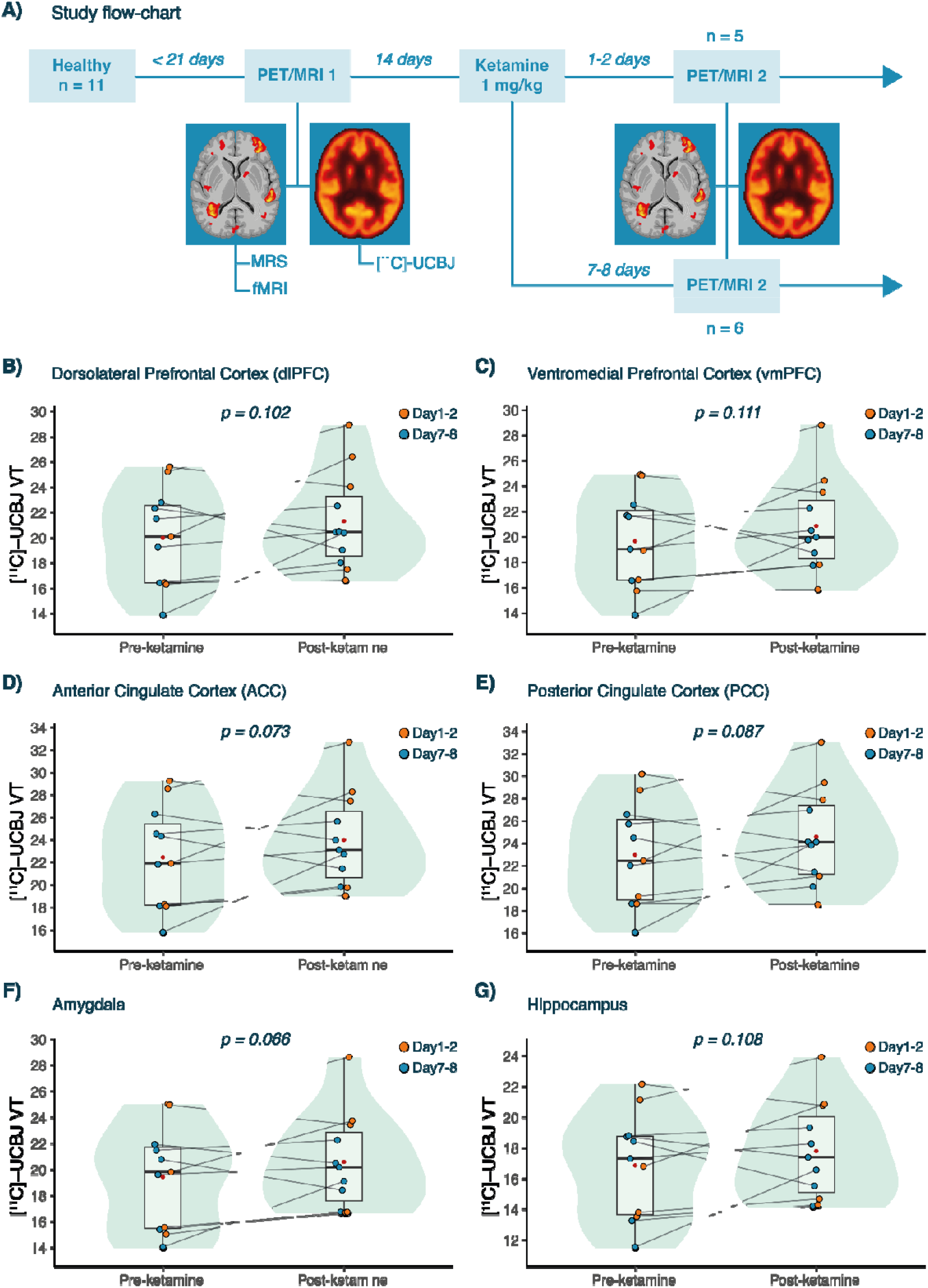
Effects of ketamine on [ C]-UCBJ VT. A) The flowchart of the study design. B-G) The change in [ C]-UCBJ VT in the B) dlPFC, C) vmPFC, D) ACC, E) PCC, F) Amygdala, and G) Hippocampus.

### 1.4. PET/MRI acquisition and processing

The PET data were acquired using an integrated General Electric Signa 3 Tesla combined PET/MRI scanner with a 32-channel head coil at the Invicro Imaging Centre (now ‘Perceptive’), London, UK. The [^11^C]-UCBJ tracer was synthesized onsite and administered i.v. as a bolus over 20 seconds by the study physician. The PET scan acquisition time was 90 minutes in addition to 30 minutes of MRI scanning. For the PET modelling, the study employed the 1 tissue compartment model for reversible binding to correlate the parent plasma input function with tissue time-activity curves, producing estimates of the total VT for each predefined ROI. The fMRI data were acquired using an Echo-Planar Imaging sequence sensitive to BOLD contrast with the following settings: TR = 2000, TE = 30, flip angle = 80°, 3mm x 3mm in-plane resolution (64 x 64 matrix), slice thickness = 3.6 mm, 36 axial slices, 240 volumes, 10 minutes. In addition a T1 weighted IR-SPGR sequence was acquired to provide an anatomical image (TI = 400 ms, TE = minimum, flip angle = 11°, 256 x 256 matrix, 1 mm isotropic voxels, sagittal slices). MRS data included two ^1^H-MRS scans: a PRESS-PROBE sequence (TE=30ms, 2×2×2cm3, 96 averages) and a GABA- and GSH-edited HERMES sequence (TE=76ms, 2.5×2.5×3cm, NEX=8). Voxels were positioned in the ACC. After quality control, datasets from 2 subjects on the PRESS data were excluded, and 7 time-points were excluded on the HERMES data. Thus, the data from the HERMES sequence were excluded from the main analysis (Supplementary Table 1). See Supplementary Data for full details on PET/MRI acquisition and processing.

### 1.5. PET analysis

Pre-defined ROIs were the dorsolateral (dlPFC) and ventromedial (vmPFC) pre-frontal cortex, ACC, PCC, hippocampus, and amygdala. The selection of the ROIs was based on previously published work on [^11^C]-UCBJ VT changes induced by ketamine [^14^]. For additional analysis across regions see Supplementary Data. The ROIs-specific VTs, corrected for subregional volume, were the primary outcome measures of the study. Additional PET metrics were the Distribution Volume Ratio-1 (DVR-1), computed by dividing the ROI VT by the centrum semiovale (CS) VT (reference region) minus 1, and the free-fraction corrected [^11^]UCB-J (FP) (See Supplementary Data). For all analyses involving the combination of PET data with fMRI metrics, [11C]-UCBJ VT was selected as the main PET outcome. This choice was motivated by several reasons. First, VT was our pre-specified primary outcome measure. Also, there was a trend in the data suggesting a difference in the control region VT between pre- and post-ketamine visits, which could potentially influence DVR-1 estimates. Lastly, no significant correlations were observed between the change in VT and the FP and DVR-1 metrics (Supplementary Data). Therefore, a systemic effect of ketamine on tracer metabolism may have introduced variance into the free-fraction or reference region-corrected metrics.

### 1.6. fMRI Amplitude of low frequency fluctuations (ALFF)

To compute whole-brain intrinsic activity, ALFF measures were calculated using the Analysis of Functional Neuroimages (ANFI v.20.1.06) 3dRSFC module. ALFF is the power of the BOLD signal in the 0.01-0.1 Hz low frequency range. The data in these analyses were band-pass filtered using a range of 0.01-0.1Hz. As ALFF is sensitive to the raw values of the BOLD time series (which are arbitrary values) some normalisation of ALFF measures is standard practice [^38^]. In this case, Z-normalisation was used after ALFF analysis, where the mean of each participant’s ALFF image was divided by its standard deviation, to give an overall Z-score brain map for each participant (Supplementary Data).

### 1.7. fMRI mutual information functional connectivity (miFC)

Whole-brain intrinsic FC was computed as the pairwise mutual information (miFC) between regional timeseries in line with previous work [^39^]. Brain regions were defined using the 400-region Schaefer atlas, which assigns each region to a resting state functional network [^40^]. Regional timeseries were computed as the average of all voxels within a region, and were Z-scored to centre mean 0. Pairwise miFC was computed using the algorithm from Peng et al [^41^].

### 1.8. fMRI Graph theoretic metrics

Whole-brain graph theoretic metrics, such as brain modularity and global connectivity, can be extrapolated from functional connectivity metrics to assess how information is communicated across the brain. Modularity assesses whether a graph exhibits small-world architecture through the presence of modules [^42^], while global brain connectivity is a measure of the degree of interconnectedness of each graph node with all the others. We computed the modularity of the miFC connectivity matrix for each subject and session using the Brain Connectivity Toolbox (BCT) using a similar approach to [^43^]. Global brain connectivity was assessed by computing the average miFC of each voxel with all other grey matter voxels in the brain, similarly to [^18^].

### 1.9. Multimodal statistical analysis

A one-sided linear mixed-effect model was used for within-subject analysis measuring changes in plasma free-fraction, VT, DVR-1, and FP measures of [^11^C]-UCBJ before and after ketamine administration (e.g., ROI_VT ∼ Time + (1|Subject)). See Supplementary Data for analysis of ketamine effects across multiple regions. An interaction term was added to the model to test for the between-subject difference in [^11^C]-UCBJ metrics change before and after ketamine administration between participants who had their Scan 2 at 1 to 2 days following ketamine and those who had it at 7 to 8 days. For all models, the effect size was estimated using Cohen’s d. Also, the confidence of the result was estimated by computing the Bayesian Factor (BF). The same statistical approach was adopted to test pre-to post-ketamine differences in ACC Glutamate concentrations (Two-sided). Pair-wise Spearman’s correlation tests were used to analyse the correlation between [^11^C]-UCBJ VT, BP, and FP metrics and their difference following ketamine. Similarly, Spearman’s correlation test was used to test for the correlation between glutamate concentrations and [^11^C]-UCBJ within the ACC. To account for FDR inflation due to multiple comparisons, the p-values resulting from Spearman’s tests were adjusted independently using the Benjamini-Hochberg adjustment. To measure the effects of ketamine on ALFF, we implemented a voxel-wise paired analysis using FSL’s FEAT with a second-level mixed-effects model (FLAME1). Results were thresholded at Z=3.1, with a cluster p<0.05.

To test for FC difference following ketamine administration, permutation tests evaluated the effect of the session on pairwise miFC, modularity, and global connectivity (FDR corrected; See Supplementary Data).

To explore the correlation between the structural and functional measures, acquired with PET and fMRI respectively, a whole-brain data-driven approach was adopted. Voxel-wise Pearson’s correlations were performed between Δ[11C]-UCBJ VT and ΔALFF datasets, following methods in [^44,45^], to explore correlations between PET and fMRI metrics in a whole-brain, data-driven approach. To further explore the informational role of those brain regions exhibiting significant voxels of Δ [^11^]UCB-J VT/Δ ALFF correlations (see Results), the graph-theoretic metric of betweenness centrality (BC) was extrapolated from the pairwise miFC for those regions for each subject and session [^46^]. Statistical analyses were conducted in MATLAB 2023A and R 2024.

## 2. Results

### 2.1. Effects of ketamine on [^11^C]-UCBJ

Assessments of the normality and quality of the PET data is reported in Supplementary Data. With all subjects included in the analysis (n = 11), we observed no statistically significant increase in VT of [^11^C]-UCBJ following ketamine administration in any of the analysed ROIs (Supplementary Table X). Notably, there was a trend towards an increase in all ROIs, approaching statistical significance in the amygdala (Figure 1F; M: 7.33 ± 13.44 %; β = 1.17; p = 0.066; d = 0.495; BF = 0.837) and the ACC (Figure 1D; M: 8.53 ± 16.38 %; β = 1.54; p = 0.073; d =0.475; BF= 0.778). Similar results were obtained when the [^11^C]-UCBJ VT was corrected for plasma free-fraction to obtain FP, and for VT within the reference region CS to obtain DVR-1. However, for FP and DVR-1 no trends were observed in any ROI (Supplementary Table 2). At baseline, the VT and FP [^11^C]-UCBJ PET metrics were positively correlated in all analysed ROIs. No correlation was found between [^11^C]-UCBJ DVR-1 and the other PET metrics (Supplementary Table 5). The post-ketamine changes in [^11^C]-UCBJ VT, DVR-1, and FP were not correlated between them (Supplementary Table 6). Overall, no significant difference in the magnitude and direction of [^11^C]-UCBJ change following ketamine was detected between Day 1-2 and Day 7-8 groups, except for the amygdala (β =- 0.28; p = 0.023, d = 1.653): in this region, the Day 1-2 group showed a higher increase in [^11^C]-UCBJ DVR-1 as compared to the Day 7-8 group. All subsequent analysis are shown for the full sample as no significant difference in the effect of ketamine on our primary outcome [^11^C]-UCBJ VT was detected between the Day 1-2 and Day 7-8 groups and to preserve statistical power.

### 2.2. Effects of ketamine on ACC glutamate

Ketamine produced a statistically significant increase in glutamate levels within the ACC (Figure 2A; n = 9; M: 21.77 ± 25.35 %; β = 2.92; p = 0.030, d =0.878; BF =2.70). There was a difference between the Day1-2 and Day7-8 in the magnitude of the pre- to post-ketamine change in ACC glutamate levels such that the Day7-8 group showed greater increases as compared to the Day 1-2 group (β = -4.32; p = 0.042: d =1.662). See Supplementary Data for other MRS metrics.

**Figure 2.**
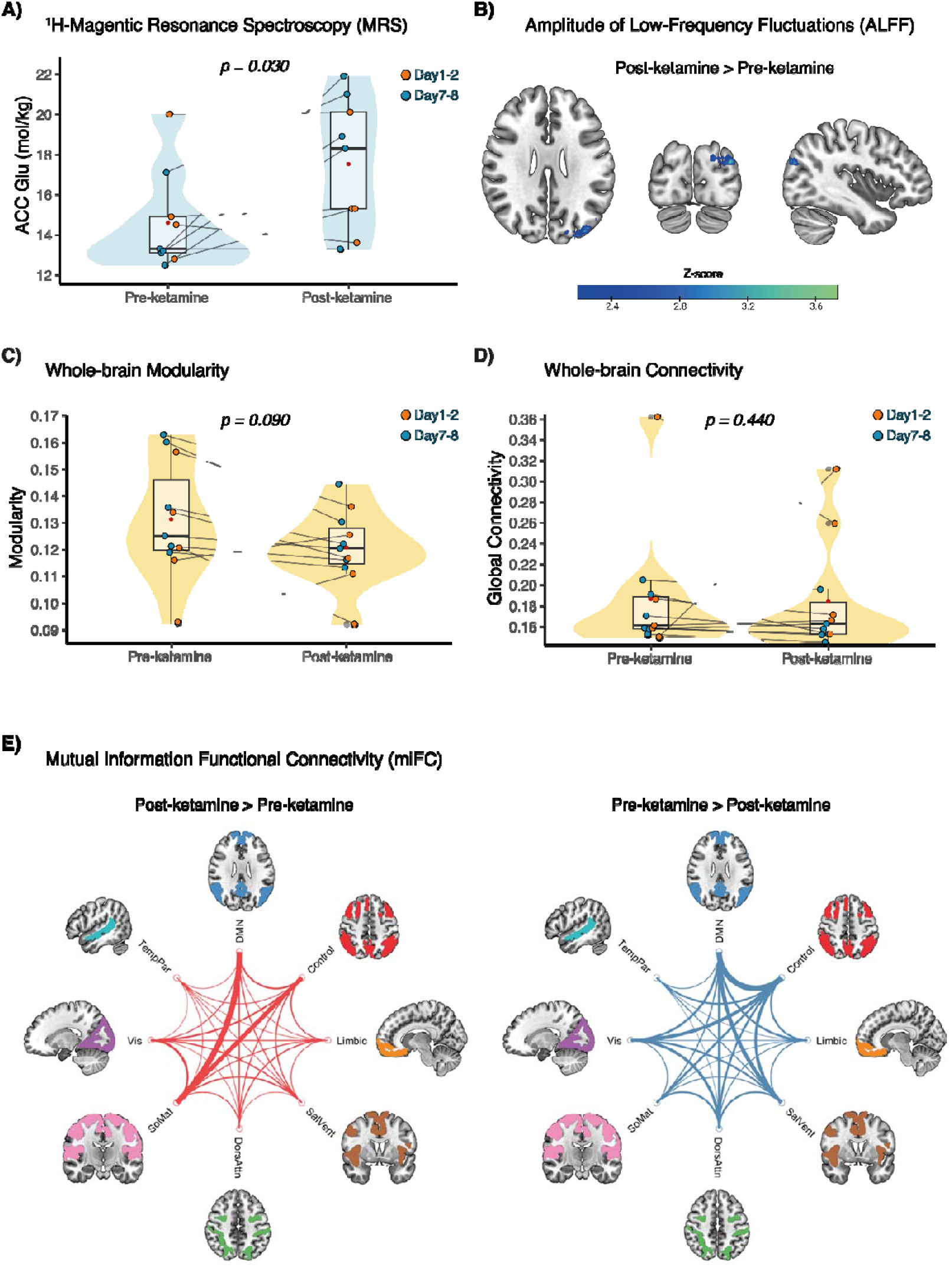
Effects of ketamine on MRI measures. A) Change in ^1^H-MRS Glutamate in the ACC. B) Change in whole-brain ALFF. C) The change in whole-brain modularity and D) global connectivity. E) Circle plots show edges with increased (red) or decreased (blue) in miFC from session 1 to session 2. All regions are grouped into a functional network, and differences in miFC between regions in different functional networks are shown with lines between the networks. The thickness of the line scales with the number of regions showing significant differences in miFC. Abbreviations are as follows: SalVent=Salience/Ventral Attention; DorsAttn=Dorsal Attention; SoMat=Somatomotor; Vis=Visual; TempPar=Temporal Parietal.

### 2.3. Effects of ketamine on whole-brain ALFF, miFC, modularity, and global connectivity

Regional ALFF was significantly lower post-ketamine only in a small region of the occipital cortex (n= 11; FLAME-1, Z>3.1, cluster p<0.05; Figure 2B). Overall, most of the statistically significant changes (71%) in miFC after dosing were decreases (n=11; Supplementary Table 8). Edges with significantly decreased post-dose miFC were primarily inter-vs intranetwork (83% vs 17%, respectively). Inter-network changes in miFC included the DMN-Control network, and from both the DMN and Control to Salience/Ventral Attention or Visual network (Figure 2E). Edges with increased miFC from pre- to post-dose were also primarily inter-vs intranetwork (92% vs 8%, respectively). The networks showing increased miFC post-dose were concentrated between the DMN and Control network to the Somatomotor network (Figure 2E).

Ketamine-induced differences in resting-state whole-brain network dynamics were evaluated through estimates of modularity and global connectivity extrapolated from miFC. There was a trend towards a statistically significant decrease in modularity from pre- to post-dose (t-stat=0.01, p=0.09, d=0.60; Figure 2C). There was not a significant difference in global connectivity between sessions (t-stat=0.002, p=0.44, d=0.04; Figure 2D).

### 2.4. Correlation between ACC [^11^C]-UCBJ VT and glutamate

No statistically significant relationship was found between changes in glutamate and [^11^C]-UCBJ VT within the ACC following ketamine administration (n=9; ρ = 0.01, p = 0.980).

### 2.5. Correlation between whole-brain [^11^C]-UCBJ VT, ALFF, and BC

There were significant negative correlations between ΔALFF and Δ[^11^C]-UCBJ VT suggesting that a greater positive change in VT was associated with decreased ALFF following ketamine administration. As reported in Figure 3A, these correlations were found in regions spanning the PCC, precuneus, parietal regions, precentral gyrus, postcentral gyrus, superior frontal gyrus, right temporal lobe, overlapping with DMN regions (n=11; Supplementary Table X). A significant negative relationship between higher post-dose [^11^C]-UCBJ VT and lower post-dose BC was observed in the PCC (n=11; r=-0.69, p=0.02; Figure 3B). There were no significant relationships between changes in [^11^C]-UCBJ VT and BC in the other tested regions (Supplementary table X).

**Figure 3.**
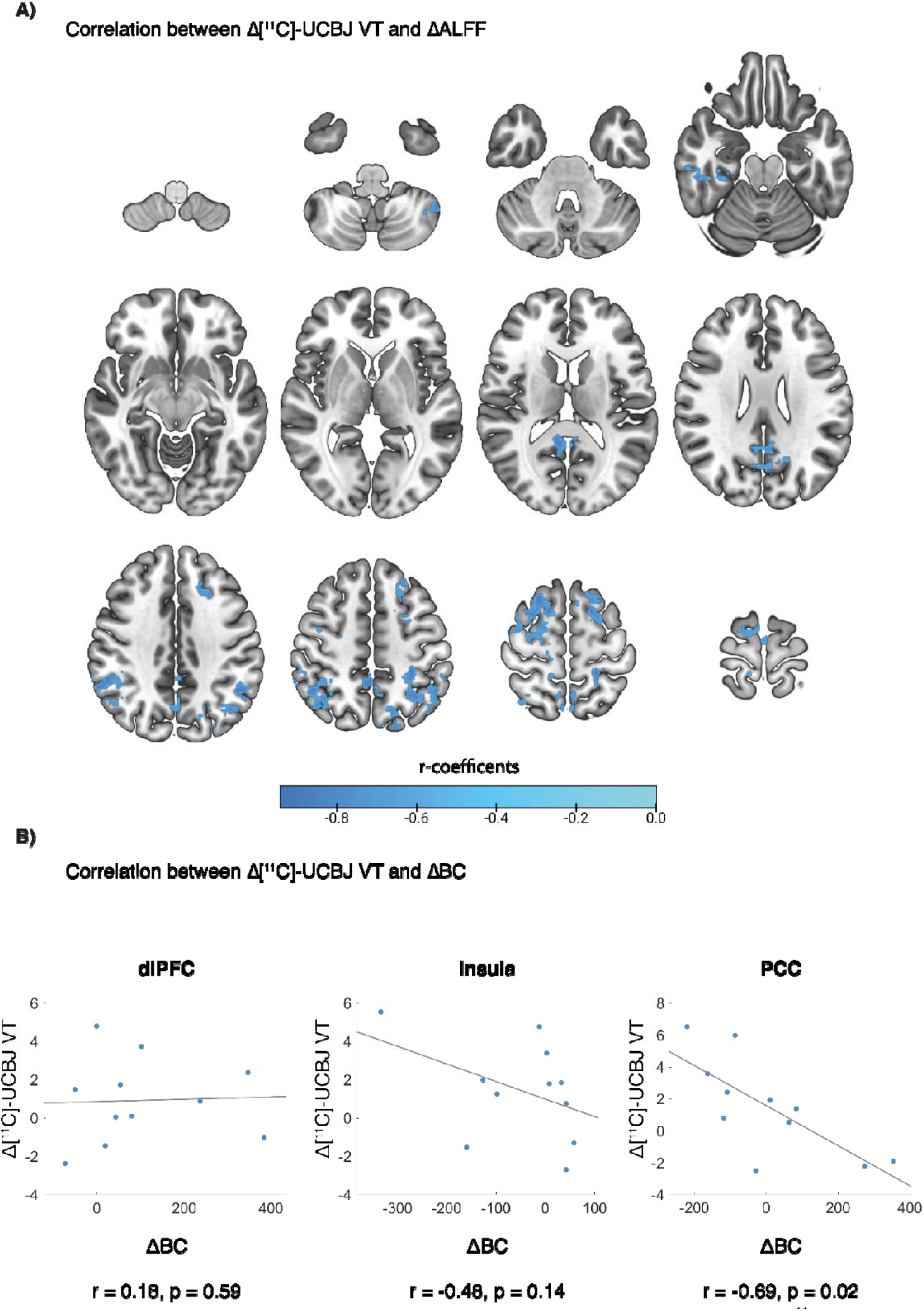
Correlation between PET and MRI measures. A) Correlation between whole-brain Δ[ ^11^C]-UCBJ VT and ΔALFF. Correlation between Δ[ ^11^C]-UCBJ VT and Δ in the dlPFC, Insula, and PCC.

## 3. Discussion

Our multimodal PET/fMRI analysis revealed that ketamine-induced alterations in synaptic plasticity were inversely related to changes in intrinsic regional brain activity, particularly in areas belonging to the DMN. Notably, the PCC emerged as a pivotal region linking the effects of ketamine at the molecular level with whole-brain dynamics, evidenced by a significant negative association between changes in synaptic plasticity and the PCC’s network influence in information processing, as measured by betweenness centrality. Taken together, our findings suggest a complex enduring effect of ketamine on neural plasticity across different scales, from synapses to high-order functional networks.

There was a trend towards an increase in [^11^C]-UCBJ VT following ketamine in our pre-defined ROIs, however, the change was not statistically significant at the group-level, and the effect size and confidence of the result were relatively weak. The trend was further explored with an additional analysis across the pre-defined ROIs and whole-cortex (See Supplementary Data). When analysing all pre-defined ROIs collectively in a single model, we observed a statistically significant increase in [^11^C]-UCBJ VT (p < 0.001). The whole-cortex analysis yielded similar significant results (p < 0.001). However, when we calculated volume-weighted averages of either the pre-defined ROIs or all cortical regions, the results mirrored our initial findings, showing a non-significant trend (p = 0.092). An in depth discussion of these analysis is presented in the Supplementary Data. We did not observe differences between participants scanned at 1-2 days following ketamine and those scanned at 7-8 days, in any of the analysed PET metrics and ROIs. The only exception was in the amygdala, where we detected a significant increase in [^11^C]-UCBJ DVR-1 in the Day1-2 specifically. Although this finding showed a considerable effect size, we also observed a trend suggesting a differential binding of [^11^C]-UCBJ VT in our reference region (CS) following ketamine. This could signify a systemic effect of ketamine on [^11^C]-UCBJ metabolism. Also, while [^11^C]-UCBJ VT and FP values were positively related at baseline, as expected, their change following ketamine did not show a linear relationship, reinforcing the possibility of a systemic effect of ketamine on the tracer. Overall, our results do not provide conclusive evidence of a group-level increase in synaptic plasticity induced by a single psychedelic dose of ketamine (1 mg/kg) in healthy human brains at 1 to 8 days post-administration, suggesting high individual variability in drug neuroplastic response. This broadly aligns with the study by Holmes et al. (2022), which also provided inconclusive evidence regarding a unidirectional effect of ketamine (0.5 mg/kg) on synaptic SV2A, as measured via [^11^]-UCBJ, in both healthy and depressed individuals 1 day post-drug administration [^14^]. Several factors could account for the absence of sizable changes in [^11^C]-UCBJ following ketamine. While we employed double the dosage of Holmes et al. (2022), approximating the highest titration ketamine dose used in psychiatric practice [^34^], the absence of a robust effect on [^11^C]-UCBJ suggests that changes in synaptic plasticity in humans (if it could be imaged at all) might require multiple doses to be detected via PET. This is supported by research showing that some patients require multiple ketamine doses to achieve therapeutic benefit, associated with significant structural and functional alterations [^47–49^]. Another possibility is that ketamine’s effects on plasticity may be more pronounced in depression patients, as supported by some clinical [^14,49^] and preclinical observations [^50,51^]. Also, [^11^C]-UCBJ binding reductions were detected in advanced [^12^] but not early-stage schizophrenia [^52^], suggesting that [^11^C]-UCBJ may be more effective as a biomarker in cases of substantial neural remodelling, such as observed in severe patient populations. Lastly, multiple physiological factors may have influenced the [^11^C]-UCB-J VT results (see Supplementary Data for discussion on sleep). Despite these considerations, the presence of a trend in our data suggests a potential biological effect that warrants further investigation in larger cohorts.

Within the ACC, ketamine produced a statistically significant increase in glutamate concentrations, with the effect particularly pronounced in the Day 7-8 group. The glutamate ‘surge’ induced by psychedelic doses of ketamine is generally thought to be acute [^53,54^], not extending beyond the drug’s presence in the body as previous studies did not detect significant changes in glutamate concentrations at 1-2 days post-ketamine in either healthy subjects or depressed patients [^55,56^]. However, using high-resolution MRS, Li et al. (2017) found an increased glutamine/glutamate ratio measured in the ACC 1 day post ketamine administration in healthy volunteers [^57^], while another study found a significant glutamate increase at 7 days [^58^]. Alongside our findings, the persistence of elevated glutamate from 1 to 8 days post-administration challenges the model of ketamine-induced glutamate increase as purely transient. Differences in scanning sequences and acquisition methodologies across studies might account for the variability in the results. Also, given our study’s limited statistical power, these results require further validation.

At the whole-brain level, we observed a trend towards a decrease in intrinsic brain activity, measured via ALFF, which was statistically significant only in a small region of the occipital cortex. Previous studies using fractional ALFF, instead of raw ALFF, during ketamine infusion reported either global [^18,59^] or occipital-specific [^19,60^] reductions, aligning with our observation. Notably, one of these studies indicated that the effect diminished by 1 day post-administration, which may account for the lack of statistical significance observed in our data [^19^]. ALFF quantifies the power of infra-slow (0.01-0.1 Hz) BOLD signal fluctuations at the voxel level. Multimodal evidence indicates BOLD signal changes correlate with high-frequency synchronized neural activity, as measured via integrated fMRI/electroencephalography (EEG). In landmark studies, it was shown that regional ketamine-induced BOLD signal changes and connectivity patterns follow different temporal profiles of EEG desynchronization of oscillatory activity [^21,59,61^]. However, no clear relationships between the EEG metrics and fractional ALFF were detected during ketamine administration, suggesting a complex relationship between those metrics under pharmacological perturbation [^59^]. While the reduction of ALFF observed in our study might be a product of ketamine-induced perturbation of synchronous neural activity in the infra-slow range or at higher frequencies, attenuated in effect size due to the temporal distance from the infusion, more research is needed to elucidate the physiological source of this effect.

We observed a significant alteration in intrinsic connectivity within and across canonical functional networks, accompanied by a trend towards reduced brain modularity. The observed reduction in network integrity and inter-network communication, particularly between high-order networks, alongside increased connectivity between low- and high-order networks, aligns with findings from previous studies on ketamine and classic psychedelics [^16,17^]. A working framework suggests that flattening of the cortical hierarchy is a hallmark of the acute effects of altered states of consciousness induced by compounds like ketamine and classic psychedelics. Compelling evidence has shown that these compounds’ effects on whole-brain functional dynamics are persistent, lasting from 1 to up to 10 days post-ketamine [^19,24,25,27,28,62^] and up to 21 days post-classic psychedelics [^43,63^]. Our findings similarly suggest that ketamine’s acute effects on functional dynamics may endure for several days after a single administration, as hypothesized.

Our data-driven multimodal analysis revealed intriguing neural dynamics associated with ketamine-induced alterations of synaptic plasticity. Firstly, a negative relationship was observed in DMN regions between [^11^C]-UCBJ VT and ALFF, meaning that participants showing an increase in synaptic plasticity following ketamine also showed reductions in intrinsic brain activity in the DMN. Further, a negative correlation between [^11^C]-UCBJ VT and BC specifically in the PCC region of the DMN, signifying a reduced role of the PCC in mediating information flow in the brain of those subjects with increased synaptic plasticity in that region. It has been proposed that the DMN involvement with narrative-self-referential processes makes it responsible, when overactivated, for ruminative and negative thought loops, characteristics of depressive symptomatology [^22,23^]. In fact, the DMN regions, especially its posterior hub, are theorized to serve as an orchestrator of information flow and gateways of directed information transfer across the brain [^64^]. Luppi et al. provided converging evidence on the role of posterior DMN as a central hub of synergistic interactions with the rest of the brain and the mediator of cognitive state transitions, which are acutely disrupted by pharmacological alterations of consciousness [^65,66^]. Hence, similar to classic psychedelics [^67^], the perturbation of DMN functioning produced by ketamine may re-calibrate the network activity by flattening its apical position in the cortical hierarchy and promoting its integration with bottom-up processes. Our results also suggests that ketamine’s modulation of DMN activity is sustained in the long-term and is paralleled by increases in synaptic plasticity. We speculate that the observed effect of ketamine may offer a precise spatiotemporal window for therapeutic intervention. However, our interpretation is tentative and specific hypothesis-driven approaches are required to confirm the validity of our findings.

Importantly, our experimental design presents several limitations. First, the limited sample size (n = 11) significantly limited our statistical power. Additionally, participants were scanned at varying time points post-ketamine (1 to 8 days) to capture a comprehensive view of ketamine’s temporal effects on [^11^C]-UCBJ. However, this approach reduced the specificity of our group-level analyses. Limitations inherent to the [^11^C]-UCBJ tracer also present challenges for examining the subtle and dynamic neuroplastic effects induced by pharmacological interventions in humans *in vivo*. First, [^11^C]-UCBJ targets a protein that is ubiquitously expressed across neuronal populations, failing to capture the neural population-specific effects captured by animal research [^3^]. Indeed, it is likely that the spatial and temporal resolution of [^11^C]-UCBJ PET imaging is insufficient to accurately capture the dynamic synaptic alterations induced by ketamine in the living brain. Further, most of the available pre-clinical evidence indicates ketamine and classic psychedelics primarily exert post-synaptic effects on neuroplasticity, such as increased dendritic spine density, with only a fraction of these spines maturing into functional synapses. Given that the binding site of [^11^C]-UCBJ is SV2A, which is located on neurotransmitter-carrying vesicles at the axon terminal, the marker may be inherently constrained in detecting ketamine-induced synaptic effects [^10^]. Moreover, studies have reported ketamine’s inhibitory effects on neurotransmitter recycling and vesicle trafficking, adding further complexity to interpreting SV2A as a plasticity marker [^68^]. Lastly, increases in SV2A could also result from a rise in the number of vesicles at the presynaptic terminal, potentially due to Hebbian learning, or from changes in the number of SV2A molecules per vesicle [^69^].

Therefore, while [^11^C]-UCBJ is among the most precise tracers currently available for in vivo imaging of synaptic plasticity, the specific process it captures remains uncertain; thus, that our results reflect structural neuroplasticity is an unverified assumption.

Despite these limitations, our comprehensive multimodal assessment —from cellular to whole-brain characterization— yielded promising results, serving as a proof-of-concept for using integrated PET/MRI to characterize ketamine’s pharmacodynamic effects. While unimodal effect sizes for the main effects of ketamine on PET, fMRI, and MRS metrics were limited, the synergy across modalities provided novel insights. Our data-driven approach highlighted the DMN regions as potential substrates for ketamine’s cross-modal effects, linking changes in a synaptic plasticity marker with intrinsic brain activity and whole-brain dynamics. Specifically, the PCC emerged as a central hub in the reorganization of functional brain hierarchies across scales [^70^], offering valuable guidance for future hypotheses with meaningful clinical potential in developing targeted interventions.

## 4. Data availability

Data and codes are available upon direct request to the authors. Address correspondence to Claudio Agnorelli at for data availability. Address correspondence to Danielle Kurtin at for fMRI analysis code availability. Address correspondence to Ekaterina Shatalina at for PET/MRI analysis code availability.

## 5. Author contributions

C.A. contributed to study design and setup, data collection, statistical data analysis, and manuscript writing. J.P. contributed to the study design and setup, data collection, and manuscript writing. G.S. contributed to the data collection and manuscript writing. D.K. contributed to the PET/fMRI data analysis, figures, and manuscript writing. E.S. contributed to the PET/fMRI data analysis and manuscript writing. K.A. contributed to the study design and setup, data collection, and manuscript writing. M.W. contributed to study design and setup, the PET/fMRI data analysis, and manuscript writing. C.R. contributed to MRS data analysis, and manuscript writing. K.G. contributed to study setup, the PET/fMRI data analysis, and manuscript writing. N.E. contributed to fMRI data analysis, and manuscript writing. G.S. contributed to PET analysis and manuscript writing. K.Z. contributed to the data collection and manuscript writing. B.W. contributed to statistical data analysis and manuscript writing. A.F. contributed to study supervision and manuscript writing. I.R. contributed to study design and setup, study supervision, statistical data analysis, and manuscript writing. R.C-H. contributed to study design and manuscript writing. D.N. and P.M. contributed to acquisition of funding, study design and setup, as well as manuscript writing. D.E. contributed to study design and setup, study management, data collection, study supervision, and manuscript writing.

## 6. Funding

GE Healthcare Global Research Organization’s Investigator-Initiated Study Program

7. Acknowledgements

We thank GE for funding support and Imperial College London for sponsorship and study governance. A special thanks to Central & North West London NHS Foundation Trust and its staff for supporting and hosting the study in the CNWL-Imperial collaborative research space, CIPPRes Clinic at St Charles Hospital. We thank the NOCLOR team for CIPPRes-site governance, CNWL’s lead pharmacists Tf Chan and Allan Sebti for invaluable help with study drug processes, as well as Trust Directors Drs Jo Emmanual, Jasna Munjiza, and Cornelius Kelly for welcoming and hosting the Imperial-based research team in CIPPRes Clinic. We thank Martin Osugo for help with study procedures. We also thank all the volunteers for their participation.

