## Supplementary Data for "Detecting neuroplastic effects induced by ketamine in healthy human subjects: a multimodal approach"

1. Participants screening

Participants (n=100) were recruited through word of mouth and public advertisements. Initial telephone screening assessed eligibility based on predetermined criteria: age between 20-60 years, absence of physical or psychiatric conditions, and no history of substance abuse. Recent use (within 6 months) of ketamine or classic psychedelics was exclusionary. Following screening, 30 candidates underwent comprehensive in-person physical and psychological evaluations. These assessments, conducted by the study physician, included informed consent, electrocardiogram (ECG), blood pressure measurement, blood screening, alcohol breathalyser, and urine drug testing, along with detailed substance use history assessment. Eleven healthy male participants (mean age 32 ± 10 years) met all inclusion criteria and were enrolled in the study. While all participants had prior experience with classic psychedelics, seven were ketamine-naïve. Participants were required to maintain abstinence from alcohol and illicit substances for one week prior to and throughout the study duration. All procedures were conducted at the St Charles Centre for Health & Wellbeing Hospital (London, UK) in accordance with institutional guidelines and received ethical approval from the Brent Research Ethics Committee, London, UK.

2. Drug administration setting

Racemic ketamine (1:1 enantiomeric mixture) was administered intravenously at 1 mg/kg over 40 minutes under continuous ECG and blood pressure monitoring. The drug was sourced from the St Charles Centre for Health & Wellbeing Pharmacy and administered by the study physician at the institution's CIPPRes Clinic. Durg administration occurred in an enriched setting inspired by psychedelic-assisted psychotherapy protocols [^1^]. The treatment room featured ambient lighting, plants, and art installations. Participants were provided with blindfolds and exposed to a customized musical composition designed specifically for the drug-induced experience. The study physician and a researcher maintained continuous presence throughout each infusion session. No serious adverse events were reported

3. PET acquisition and processing

The PET data were processed with the following steps. Continuous blood data were acquired through arterial cannulation and calibrated to align with overlapping discrete whole blood samples, forming a comprehensive whole blood activity curve that spanned the 90-minutes scan duration. Activity measurements from the discrete plasma samples were juxtaposed with the corresponding whole blood data, forming the plasma-over-blood (POB) data. This was then fit to a constant model, enabling interpolation of the relationship. The POB model fit curve was multiplied by the whole blood curve, culminating in an estimated total plasma curve. Plasma samples underwent HPLC analysis to determine the parent fraction, which was subsequently fit to a sigmoidal model. The parent plasma input function, derived by multiplying the ensuing parent fraction profile with the total plasma curve, was further refined by smoothing it post-peak using a multi-exponential fit. An incorporated time delay was adjusted for in the kinetic modelling. All image data were analysed using Invicro London in-house PET data quantification tool, MIAKATTM. MIAKATTM is implemented using Matlab and makes use of SPM12 functions for image segmentation and registration. Each participant's MRI image underwent grey matter segmentation and was then registered to an anatomical template image in MNI152. Dynamic PET images, registered to the MRI scans of participants, were corrected for any motion. An automated definition of ROIs was performed on the MNI152 space based on the CIC atlas [^2^]. The ROIs, defined on the MRI images, were applied to the dynamic PET data to extract regional time-activity curves. For the PET modelling, the study employed the 1 tissue compartment model for reversible binding to correlate the parent plasma input function with tissue time-activity curves, producing estimates of the total VT for each predefined ROI. Pre-defined ROIs were the dorsolateral (dlPFC) and ventromedial (vmPFC) pre-frontal cortex, ACC, PCC, hippocampus, and amygdala. Those ROIs were extracted from the CIC atlas and defined as follows: dlPFC included the anterior and posterior dlPFC, the vmPFC included anterior and posterior vmPFC, the ACC included the ventral cingulate subcallosal gyrus, the anterior cingulate gyrus, and the dorsal anterior cingulate grey matter. The PCC, amygdala, and the hippocampus were taken directly from the CIC atlas. The selection of the ROIs was based on previously published work on [^11^C]-UCBJ VT changes induced by ketamine [^3^]. The ROIs-specific VTs, corrected for subregional volume, were the primary outcome measures of the study (i.e., synaptic density). Additional PET metrics were the DVR-1 and the FP (See Materials and Methods). The FP is obtained by dividing the unspecific binding of [^11^]UCB-J in the plasma from the ROI-specific VTs, as in Equation 1. The DVR-1 of each ROI was obtained by dividing the ROI-specific VTs by the VT of the reference region, the centrum semiovale (CS), according to Equation 2.

*Equation 1: [^11^]UCB-J Free fraction correction*

$\text{FP= }\frac{\text{VT (ROI)}}{\text{VT(Plasma)}}\text{ }$

*Equation 2: [^11^]UCB-J Reference region correction*

$\text{DVR - 1}\text{= }\frac{\text{VT (ROI)}}{\text{VT(CS)}}\text{-1}$

4. MRS data acquisition and processing

MRS included two ^1^H-MRS scans: a PRESS-PROBE sequence (TE=30ms, 2x2x2cm3, 96 averages) and a GABA- and GSH-edited HERMES sequence (TE=76ms, 2.5x2.5x3cm, NEX=8). Voxels were positioned in the ACC. Non-water suppressed spectra were acquired for water scaling and eddy-current correction. Spectra from acquisitions were pre-processed, analysed and quantified with Osprey using default parameters [^4^]. Quality of the data was determined by visual inspection and determining the 99% CI of the 1H-FWHM as the linewidth upper bound. Datasets from two subjects on the press data were excluded, and 7 time-points were excluded on the HERMES data (Supplementary Table 1). Molal metabolite concentrations are reported relative to water and corrected for tissue type. Quality of the data was determined by visual inspection and determining the 99% CI of the 1H-FWHM as the linewidth upper bound. After quality control, datasets from two subjects on the PRESS data were excluded, and 7 time-points were excluded on the HERMES data. (Supplementary Table 1).

| **Subject Number** | **Time point** | **ACC PRESS data quality** | | **ACC HERMES data quality** | |
| --- | --- | --- | --- | --- | --- |
|  |  | **Creatine SNR** | **H_2_O FWHM [ppm]** | **Creatine SNR** | **H_2_O FWHM [ppm]** |
| 1 | Baseline | 29 | 0.07 | 55 | 0.13 |
| 1 | Post1 | 34 | 0.06 | 61 | 0.12 |
| 2 | Baseline | 29 | 0.07 | 55 | 0.13 |
| 2 | Post1 | 34 | 0.06 | 61 | 0.12 |
| 3 | Baseline | 41 | 0.06 | 78 | 0.06 |
| 3 | Post1 | 47 | 0.04 | 109 | 0.06 |
| 4 | Baseline | 45 | 0.06 | 127 | 0.10 |
| 4 | Post1 | 41 | 0.06 | 84 | 0.06 |
| 5 | Baseline | 44 | 0.05 | 94 | 0.06 |
| 5 | Post1 | 38 | 0.05 | 74 | 0.09 |
| 6 | Baseline | 22 | 0.10 | 46 | 0.13 |
| 6 | Post1 | 32 | 0.11 | 205 | 0.13 |
| 7 | Baseline | 31 | 0.07 | 69 | 0.11 |
| 7 | Post1 | 31 | 0.06 | 86 | 0.07 |
| 8 | Baseline | 37 | 0.05 | 58 | 0.10 |
| 8 | Post1 | 37 | 0.05 | 52 | 0.07 |
| 10 | Baseline | 22 | 0.12 | 286 | 0.15 |
| 10 | Post1 | 25 | 0.09 | 115 | 0.09 |
| 11 | Baseline | 29 | 0.07 | 57 | 0.10 |
| 11 | Post1 | 34 | 0.06 | 73 | 0.08 |
| 12 | Baseline | 31 | 0.07 | 58 | 0.10 |
| 12 | Post1 | 29 | 0.07 | 45 | 0.13 |

*Supplementary Table 1: MRS data quality: grey cells show excluded data (FWHM cut-off determined from the 99% confidence interval). For the PRESS data, data from subjects 6 and 10 (baseline and post1) were excluded as they displayed linewidths above the 0.08ppm threshold. For HERMES, data from subject 1(baseline), subject 2 (baseline) subject 6 (baseline, post1), subject 10 (baseline) and subject 12 (post1) were excluded because they showed linewidths above the 0.12ppm threshold; data from subject 10 (post1) was excluded because of visually noisy baseline and poor fit.*

5. fMRI data acquisition and processing

The fMRI data were acquired using an Echo-Planar Imaging (EPI) sequence sensitive to BOLD contrast with the following settings: TR = 2000, TE = 30, flip angle = 80°, 3mm x 3mm in-plane resolution (64 x 64 matrix), slice thickness = 3.6 mm, 36 axial slices, 240 volumes. In addition a T1 weighted IR-SPGR sequence was acquired to provide an anatomical image (TI = 400 ms, TE = minimum, flip angle = 11°, 256 x 256 matrix, 1 mm isotropic voxels, sagittal slices). All functional data and anatomical data were pre-processed with FSL (FMRIB Software Library v5.0.4; http://www.fmrib.ox.ac.uk/fsl/). BET was used for brain extraction of the anatomical data and fsl_anat was used for additional anatomical data pre-processing. Motion correction was performed with FMRIB Linear Image Registration Tool (MCFLIRT), with spatial smoothing using a Gaussian kernel of full width at half maximum (FWHM) 6mm. A two-step co-registration, first to the subject’s individual anatomical image followed by registration to an anatomical template image in standard stereotactic space (MNI152) was performed, with no temporal filtering applied to ensure the data captured the full frequency range. White matter and CSF were used as regressors of no interest in addition to a set of 24 extended head-motion regressors that were included as confounds (these included temporal derivatives and quadratic functions derived from the raw head-motion parameters).

The amplitude of low-frequency fluctuations (ALFF) was extrapolated from the raw BOLD time series from each voxel. Fractional ALFF (fALFF) is a related measure which normalises ALFF by dividing by the power in the full frequency spectrum [^5^]. While fALFF may be somewhat less influenced by physiological effects it also has significantly lower test-retest reliability [^6^]; a key consideration for the current repeated-measures design. It was therefore decided that ALFF was the most appropriate measure for this study.

Functional connectivity (FC) was computed on the average timeseries of all voxels within a region. Regional parcellation was determined with the 400-parcellation Schaefer atlas [^7^], which labels each region with the resting-state network in which each region resides [^8^]. Regional timeseries were then z-scored to centre mean 0 and ± 1 standard deviation. Then, pairwise FC was computed via mutual information (miFC) as per our previous work [^9^]. While conventional, correlation-based measures of undirected functional connectivity only assess monotonic relationships, miFC is an information theoretic measure of undirected, pairwise functional connectivity that captures both nonlinear and linear relationships between regional timeseries [^10,11^]. As an information theoretic measure of FC, miFC considered “connectivity” as the extent to which the timeseries of one region reduces the uncertainty in knowing the timeseries of another region. Our previous work provides a formal representation of how miFC computed in this work [^9^]. Whole-brain graph theoretic metrics such as brain modularity and global connectivity were extrapolated from functional connectivity metrics to assess how information is communicated across the brain. Modularity assesses whether a graph exhibits small-world architecture through the presence of modules. Modules consist of several densely interconnected nodes (i.e., brain regions), and in systems with high modularity, there are relatively few connections between nodes in different modules [^12^]. We computed the modularity of the miFC connectivity matrix for each subject and session using the Brain Connectivity Toolbox (BCT) [^13^] using a similar approach to [^14^]. Global brain connectivity is a measure of the degree of interconnectedness of each graph node with all the others and was assessed by computing the average miFC of each voxel with all other grey matter voxels in the brain, similarly to [^15,16^].

6. Statistical analysis

The normal distribution of the [^11^C]-UCBJ PET data was assessed using the Shapiro-Wilk normality test. For the pre-defined ROI analysis, a linear mixed-effect model was used for within-subject analysis measuring changes in plasma free-fraction, VT, DVR-1, and FP measures of [^11^C]-UCBJ before and after ketamine administration. The VT, DVR-1, or FP of [^11^C]-UCBJ within each ROIs was the dependent variable, the time-point was the independent variable (i.e., PET/MRI 1 and PET/MRI 2), inserted as a fixed effect, and a random intercept was added for each subject to account for the paired nature of the data (e.g., ROI_VT ~ Time + (1|Subject)). Statistical significance was considered at α values below 0.05, taken from a one-sided distribution. The choice of a one-sided statistics was justified by the a-priori definition of the ROIs. An interaction term was added to the model to test for the between-subject difference in [^11^C]-UCBJ metrics change before and after ketamine administration between participants who had their Scan 2 at 1 to 2 days following ketamine and those who had it at 7 to 8 days. For all models, the effect size was estimated using Cohen’s d [^17^]. Also, the confidence of the result was estimated by computing the Bayesian Factor (BF) [^18^]. The same statistical approach was adopted to test pre-to post-ketamine differences in ACC Glutamate and GABA concentrations, as well as to test pre-to post-ketamine differences in the weighted average of [^11^C]-UCBJ VT across ROIs (pre-defined and all cortex). In addition, another set of linear mixed-effect models was used to explore the presence of a trend across ROIs by using either the pre-defined ROIs or all cortical regions. In these models, the main effect of Time across ROIs was analysed by incorporating all regions in the model and by adding random effects for each subject and region to account for the hierarchical structure of the data. The model was specified as VT ~ Time + (1|Subject:ROI), where Time was coded as a fixed effect and subject-specific regional variations were incorporated as nested random effects.

Pair-wise Spearman’s correlation tests were used to analyse the correlation between [^11^C]-UCBJ VT, DVR-1, and FP metrics and their difference following ketamine. Similarly, Spearman’s correlation tests was used to test for the correlation between glutamate and GABA concentrations and [^11^C]-UCBJ within the ACC. To account for FDR inflation due to multiple comparisons, the p-values resulting from Spearman’s tests were adjusted independently using the Benjamini-Hochberg adjustment (Results of the FDR correction are reported as "p adj. " in the main text) [^19^].

To measure the effects of ketamine on ALFF, we implemented a voxel-wise paired analysis using FSL’s FEAT with a second-level mixed-effects model (FLAME1). Results were thresholded at Z=3.1, with a cluster p<0.05.

Permutation tests evaluated the effect of session on pairwise miFC, modularity, and global connectivity. Permutation tests were carried out in line with [^20^]. For example, permutation tests for significant between-session differences in miFC begin by computing the difference in miFC between sessions, generating a t-stat. Then a null distribution of t-stats is generated by a minimum 1,000 permutations of shuffling session labels and computing the difference between “sessions”. After 1,000 permutations a Kolgov-Smirnov tests assesses whether the null distribution is normally distributed. If not, then another 5,000 permutations are iteratively added until a null distribution that meets assumptions for normalcy are met. P-values are determined as the percent of values with the same or more extreme value on the null distribution. Significance is set at p<0.05, i.e., if there is less than 5% of all values in the null distribution with either a higher or lower value than the empirically derived test statistic. P-values for the effect of session on miFC were considered significant if p<0.05 after FDR-adjustment or Bonferroni correction for multiple comparisons, respectively.

To explore the correlation between the structural and functional measures, acquired with PET and fMRI respectively, a whole-brain data-driven approach was adopted. For each brain voxel, we extrapolated the ∆[^11^]UCB-J VT (i.e., synaptic density) and ∆ALFF (i.e., brain activity) differences between pre- and post-ketamine scans for all subjects. Whole-brain correlation analyses were carried out between ∆[^11^]UCB-J VT and ∆ALFF difference maps and were concatenated into 4D datasets using FSL’s fslmerge tool and masked using a 75% grey mask derived from the MNI152 template. AFNI’s 3dTcorrelate module was then used to run voxel-wise Pearson’s correlations between the PET and fMRI datasets. Following methods in [^21,22^], resultant r coefficient images were transformed to t-statistical images using fslmaths and these t-maps were then transformed to Z-scores using FSL’s ‘ttoz’ function. FSL’s ‘easythresh’ function was then used to threshold the images at Z=2.3, with cluster extent brain thresholding applied at p<0.05 to correct for multiple comparisons across the brain. Final figures were made by creating a binary mask of significant clusters for each analysis and using this to mask the original r-images to show the r-coefficients corresponding to the significant findings. To further explore the informational role of those brain regions exhibiting significant voxels of ∆[^11^]UCB-J VT/ALFF correlations (see Results), the graph-theoretic metric of BC was extrapolated from the pairwise miFC for those regions for each subject and session [^23^]. While modularity and global connectivity are summary measures for an entire graph (i.e., whole-brain), BC measures the importance of a node within the graph. The BC of those regions with significant structural/functional correlations(∆[^11^]UCB-J VT/∆ALFF) was correlated with ∆[^11^]UCB-J VT.

7. Ketamine effects on [^11^C]-UCBJ
   7.1. [^11^C]-UCBJ data quality

The difference between the amount of [^11^C]-UCBJ injected before and after ketamine injection was normally distributed (W = 0.91; p = 0.244). There was no statistically significant difference in the injected mass of [^11^C]-UCBJ between the pre- and post-ketamine, Mean difference (M): -3.32 ± 54.05 %; β = -0.39; p = 0.499. No difference in plasma free-fraction was detected following ketamine (M: 4.53 ± 16.44 %; β = 0.01; p = 0.347;Table 2). There was a trend towards an increase in [^11^C]-UCBJ VT (M: 7.88 ± 12.46 %; β = 0.30; p = 0.073; Table 2) within the reference region CS from pre- to post-ketamine scans, even though it was not statistically significant. The distribution of the difference in [^11^C]-UCBJ VT concentration within the ROIs was checked for normality. Data were normally distributed in all ROIs: the dlPFC (W = 0.93; p = 0.395), the vmPFC (W = 0.94; p = 0.501), the ACC (W = 0.91; p = 0.273), the PCC (W =0.98; p = 0.883), the hippocampus (W = 0.92; p = 0.293), and the amygdala (W = 0.95; p = 0.701).

7.2. Change in [^11^C]-UCBJ following ketamine

Supplementary Table 2 shows the change in [^11^C]-UCBJ VT, DVR-1, and FP following ketamine administration in all the analysed pre-defined ROIs (i.e., dlPFC, vmPFC, ACC, PCC, amygdala, and hippocampus). To explore the presence of a trend in [^11^C]-UCBJ VT (our primary outcome measure) after ketamine across our pre-defined ROIs, data were analysed following two distinct statistical approaches. Following method similar to Johansen et al. (2023), a volume weighted average of our ROIs was calculated for each participant and timepoint [^24^]. The data were normally distributed (W=0.97; p = 0.811). Similar to the individual ROIs, a non-statistically significant trend towards an increase in [^11^C]-UCBJ VT was observed (M: 8.26 ± 17.32 %; β =1.36; p= 0.092; d =0.430; BF = 0.66) following ketamine, with a moderate effect size but with Bayes Factor indicating anecdotal evidence for the null hypothesis. In a parallel analysis, we implemented a linear mixed-effects model to test the main effect of time (pre versus post ketamine administration) on [^11^C]-UCBJ VT across ROIs. The model architecture incorporated all subject (n=11) × timepoint (n=2) × ROI (n=6) observations (Tot = 132), maintaining the full granularity of the dataset while accounting for the inherent hierarchical structure through the random effects term. The data were normally distributed (W=0.98; p =0.096). This analysis yielded a statistically significant increase in [^11^C]-UCBJ VT (M: 7.96 ± 16.01 %; β = 1.30; p < 0.001; d = 0.446; BF = 42.98) across ROIs, with the Bayes Factor providing strong evidence for the alternative hypothesis. The same set of analysis was performed at the whole cortical level, including all regions belonging to the frontal, parietal, temporal, and occipital poles plus insular cortex and cingulum. Again, at the whole-cortex level the weighted average analysis was not statistically significant (M: 8.39 ± 17.34 %; β = 1.39; p= 0.092; d = 0.430; BF = 0.66), while the analysis with all regions included yielded a significant result (M: 8.44 ± 16.16%; β = 1.32; p < 0.001; d = 0.449; BF = 1.44), with the Bayes Factor providing anecdotal evidence for the alternative hypothesis. The divergent statistical significance between these approaches likely reflects the difference in the properties of the data distributions, due to their different degrees of freedom. While the averaged approach reduces the data to 22 total observations (11 subjects × 2 timepoints), the mixed-effects model utilizes all available observations while appropriately accounting for non-independence through its random effects structure. The considerable increase in observations contributes to the enhanced ability to detect consistent effects across regions. Both approaches yielded similar effect sizes and mean percentage changes, suggesting consistency in the magnitude of the underlying biological effect across analytical methods.

Taken together, these analyses indicate a trend toward ketamine-induced elevation in [^11^C]-UCBJ VT when examined across cortical regions, with this effect reaching statistical significance in the more statistically powered mixed-effects analysis incorporating individual regional data. However, interpretation of the [^11^C]-UCBJ VT findings warrants careful consideration given the absence of concordant trends in the complementary kinetic parameters FP and DVR-1. Also, the single ROI analysis (primary analysis) did not yield significant results, likely to reduced sample size. Indeed, a post-hoc power calculation showed an achieved power of 0.34 with effect size of 0.4, α = 0.05, in a sample size of 11. Based on our observed effect size, our power calculation indicates that a sample size of 40 participants would be needed to detect an effect with 0.8 power at α = 0.05.

To conclude, we believe that while the presence of a trend at the group level is noteworthy and it might reflect a true biological effect, the limits of our current sample size prevent us from drawing conclusions on the effects of ketamine on [^11^C]-UCBJ VT in healthy subjects (see main Discussion section).

| All participants (n = 11) | | | | | | | |
| --- | --- | --- | --- | --- | --- | --- | --- |
| ROIs | [^11^]-UCBJ Metric | Scan 1 (SD) | Scan 2 (SD) | % Difference (SD) | p-value | Cohen’s d | Bayes factor |
| dlPFC | VT | 19.99  (3.88) | 21.31  (3.84) | 8.25  (17.92) | 0.102 | 0.410 | 0.622 |
|  | FP | 81.91 (15.15) | 84.09 (13.69) | 4.38  (16.73) | 0.313 | 0.152 | 0.332 |
|  | DVR-1 | 3.56  (0.31) | 3.55  (0.27) | 0.11  (9.44) | 0.443 | -0.044 | 0.300 |
| vmPFC | VT | 19.65  (3.73) | 20.85  (3.69) | 7.62  (17.24) | 0.111 | 0.392 | 0.587 |
|  | FP | 80.60 (14.93) | 82.46 (14.50) | 3.82  (16.50) | 0.336 | 0.132 | 0.323 |
|  | DVR-1 | 3.49  (0.31) | 3.45  (0.24) | -0.56  (9.09) | 0.344 | -0.124 | 0.320 |
| ACC | VT | 22.45  (4.54) | 23.99  (4.21) | 8.53  (16.38) | 0.073 | 0.475 | 0.778 |
|  | FP | 91.93 (17.42) | 94.83 (16.04) | 4.79  (16.65) | 0.273 | 0.188 | 0.351 |
|  | DVR-1 | 4.12  (0.36) | 4.13  (0.31) | 0.56  (7.76) | 0.468 | 0.025 | 0.298 |
| PCC | VT | 22.97  (4.56) | 24.59  (4.36) | 8.80  (18.35) | 0.087 | 0.443 | 0.694 |
|  | FP | 94.10  (17.53) | 97.30  (17.39) | 4.87  (16.89) | 0.266 | 0.195 | 0.356 |
|  | DVR-1 | 4.24  (0.37) | 4.25  (0.31) | 0.69  (9.22) | 0.468 | 0.025 | 0.298 |
| Amygdala | VT | 19.40  (3.93) | 20.57  (3.71) | 7.33  (13.44) | 0.066 | 0.495 | 0.837 |
|  | FP | 79.45  (15.13) | 81.30  (14.16) | 4.03  (17.47) | 0.329 | 0.138 | 0.325 |
|  | DVR-1 | 3.42  (0.27) | 3.39  (0.28) | -0.61  (6.23) | 0.347 | -0.122 | 0.319 |
| Hippocampus | VT | 16.84  (3.44) | 17.79  (3.17) | 7.24  (16.12) | 0.108 | 0.399 | 0.599 |
|  | FP | 68.98 (13.28) | 70.28 (11.88) | 3.64  (16.92) | 0.363 | 0.109 | 0.314 |
|  | DVR-1 | 2.84  (0.30) | 2.80  (0.27) | -0.97  (8.35) | 0.290 | -0.173 | 0.342 |
| CS | VT | 4.37  (0.77) | 4.68  (0.73) | 7.88  (12.46) | 0.073 | 0.603 | 1.279 |
|  | FP | 17.96  (3.10) | 18.50  (2.96) | 4.53  (16.44) | 0.542 | 0.190 | 0.352 |
| Plasma | Free-fraction | 0.25  (0.02) | 0.26  (0.04) | 4.82  (15.16) | 0.347 | 0.298 | 0.445 |

*Supplementary Table 2. Changes in [^11^C]-UCBJ metrics before and after ketamine, all participants (n = 11). The table reports the average VT, DVR-1, and FP values within the pre-defined ROIs. The p-values are the result of one-sided linear mixed-effect models with α significance threshold set at 0.05. The effect size was computed via Cohen’s d test. The statistical confidence of the result was computed via BF analysis.*

7.3. Day 1-2 versus Day 7-8 groups

When considering only participants scanned at scanned 1-2 days (Supplementary Table 2) or 7-8 days (Supplementary Table 3), the difference in [^11^C]-UCBJ VT between pre- to post-ketamine was not statistically significant in any of the pre-defined ROIs. The 2 groups also did not show a differences in the magnitude of change of [^11^C]-UCBJ VT. In the Day 1-2 group only, there was a trend toward increase in the amygdala (M: 9.39 ± 9.77 %; β = 1.78; p = 0.072) but not significantly different from the Day 7-8 group. In the Day 1-2 group there was a statistically significant increase in [^11^C]-UCBJ DVR-1 in the amygdala (M: 3.71 ± 1.69 %; β = 0.13; p = 0.003) but not in the Day 7-8 group. Indeed, the two groups differed in the magnitude and direction of [^11^C]-UCBJ DVR-1 change in the amygdala (β =- 0.28; p = 0.023). There was no significant change in plasma [^11^C]-UCBJ free-fraction after ketamine in both Day 1-2 and Day 7-8 groups, and no significant differences between them. No difference in the [^11^C]-UCBJ FP within the CS was observed in the 2 groups with no difference between them. The difference in [^11^C]-UCBJ FP after ketamine was not significant in the Day 1-2 group or Day 7-8 group in the analysed ROIs and no differences in [^11^C]-UCBJ FP change were detected between Day 1-2 and Day 7-8 groups.

| Day 1-2 group (N = 5) | | | | | | | |
| --- | --- | --- | --- | --- | --- | --- | --- |
| ROIs | [^11^]-UCBJ Metric | Scan 1 (SD) | Scan 2 (SD) | % Difference (SD) | p-value | Cohen’s d | Bayes factor |
| dlPFC | VT | 20.74  (4.45) | 22.70  (5.44) | 9.62  (14.37) | 0.115 | 0.633 | 0.776 |
|  | FP | 83.88 (15.91) | 89.19 (17.04) | 8.64  (18.98) | 0.248 | 0.335 | 0.495 |
|  | BP | 3.63  (0.26) | 3.76  (0.24) | 3.74  (8.48) | 0.191 | 0.439 | 0.570 |
| vmPFC | VT | 20.19  (4.20) | 22.07  (5.25) | 9.48  (13.48) | 0.103 | 0.674 | 0.833 |
|  | FP | 81.81 (15.66) | 87.21 (19.12) | 8.53  (18.72) | 0.240 | 0.348 | 0.503 |
|  | BP | 3.51  (0.34) | 3.63  (0.22) | 3.68  (7.65) | 0.173 | 0.477 | 0.603 |
| ACC | VT | 23.21  (5.38) | 25.43  (5.87) | 10.13 (13.00) | 0.094 | 0.709 | 0.885 |
|  | FP | 93.88 (19.69) | 100.18 (19.53) | 9.46  (20.42) | 0.247 | 0.336 | 0.496 |
|  | BP | 4.17  (0.38) | 4.34  (0.25) | 4.29  (6.14) | 0.088 | 0.736 | 0.928 |
| PCC | VT | 23.84  (5.16) | 25.97  (6.00) | 9.45  (14.85) | 0.129 | 0.588 | 0.720 |
|  | FP | 96.51  (18.77) | 102.54  (21.38) | 8.58  (20.19) | 0.268 | 0.302 | 0.476 |
|  | BP | 4.33  (0.44) | 4.45  (0.30) | 3.36  (8.18) | 0.210 | 0.402 | 0.541 |
| Amygdala | VT | 20.06  (4.79) | 21.83  (5.12) | 9.39  (9.77) | 0.072 | 0.811 | 1.059 |
|  | FP | 81.06 (17.27) | 86.02 (17.25) | 9.29  (22.64) | 0.275 | 0.292 | 0.471 |
|  | BP | 3.46  (0.26) | 3.58  (0.21) | 3.71  (1.69) | 0.003 | 2.360 | 9.949 |
| Hippocampus | VT | 17.47  (3.89) | 18.89  (4.24) | 8.63  (11.18) | 0.102 | 0.679 | 0.840 |
|  | FP | 70.67 (13.94) | 74.40 (13.89) | 8.08  (20.04) | 0.289 | 0.270 | 0.460 |
|  | BP | 2.89  (0.29) | 2.97  (0.22) | 2.89  (5.19) | 0.142 | 0.552 | 0.678 |
| CS | VT | 4.48  (0.91) | 4.76  (1.06) | 6.31  (8.93) | 0.233 | 0.627 | 0.769 |
|  | FP | 18.10  (3.22) | 18.81  (3.91) | 6.24  (22.12) | 0.695 | 0.189 | 0.428 |
| Plasma | Free-fraction | 0.25  (0.02) | 0.26  (0.06) | 2.97  (19.05) | 0.721 | 0.172 | 0.422 |

*Supplementary Table 3. Changes in [^11^C]-UCBJ metrics before and 1-2 days after ketamine, Day 1-2 group (N = 5). The table report the average VT, BP, and FP values within the pre-defined ROIs. The p-values are the result of one-sided linear mixed-effect model with α significance threshold set at 0.05. The effect size was computed via Cohen’s d test. The statistical confidence of the result was computed via BF analysis.*

| Day 7-8 group (N = 6) | | | | | | | |
| --- | --- | --- | --- | --- | --- | --- | --- |
| ROIs | [^11^]-UCBJ Metric | Scan 1 (SD) | Scan 2 (SD) | % Difference (SD) | p-value | Cohen’s d | Bayes factor |
| dlPFC | VT | 19.37  (3.56) | 20.15  (1.52) | 7.10  (21.76) | 0.303 | 0.410 | 0.776 |
|  | FP | 80.27 (11.76) | 79.84  (9.72) | 0.83  (15.44) | 0.471 | -0.031 | 0.374 |
|  | DVR-1 | 3.51  (0.38) | 3.38  (0.13) | -2.92  (9.82) | 0.184 | -0.403 | 0.542 |
| vmPFC | VT | 19.20  (3.42) | 19.82  (1.55) | 6.08  (21.04) | 0.335 | 0.184 | 0.406 |
|  | FP | 79.60 (11.29) | 78.50  (9.32) | -0.11 (14.94) | 0.424 | -0.082 | 0.380 |
|  | DVR-1 | 3.47  (0.35) | 3.30  (0.12) | -4.09  (9.25) | 0.123 | -0.537 | 0.694 |
| ACC | VT | 21.81  (4.09) | 22.79  (2.01) | 7.20  (19.92) | 0.264 | 0.277 | 0.449 |
|  | FP | 90.31 (12.54) | 90.36 (12.52) | 0.91  (13.45) | 0.496 | 0.004 | 0.373 |
|  | DVR-1 | 4.07  (0.41) | 3.95  (0.23) | -2.54  (8.05) | 0.183 | -0.406 | 0.545 |
| PCC | VT | 22.24  (4.17) | 23.43  (2.38) | 8.26  (22.27) | 0.248 | 0.300 | 0.463 |
|  | FP | 92.08  (12.68) | 92.93  (13.73) | 1.77  (14.81) | 0.445 | 0.059 | 0.377 |
|  | DVR-1 | 4.17  (0.38) | 4.08  (0.21) | -1.53  (10.17) | 0.297 | -0.232 | 0.426 |
| Amygdala | VT | 18.85  (3.34) | 19.51  (1.90) | 5.60  (16.64) | 0.278 | 0.257 | 0.438 |
|  | FP | 78.11  (10.33) | 77.36 (11.03) | -0.36 (12.24) | 0.434 | -0.072 | 0.378 |
|  | DVR-1 | 3.39  (0.32) | 3.23  (0.22) | -4.20  (6.42) | 0.075 | -0.692 | 0.956 |
| Hippocampus | VT | 16.33  (3.15) | 16.87  (1.86) | 6.07  (20.39) | 0.321 | 0.202 | 0.413 |
|  | FP | 67.58  (10.07) | 66.84  (9.84) | -0.06 (14.67) | 0.437 | -0.069 | 0.378 |
|  | DVR-1 | 2.79  (0.34) | 2.66  (0.23) | -4.19  (9.51) | 0.118 | -0.551 | 0.712 |
| CS | VT | 4.29  (0.67) | 4.61  (0.39) | 9.19  (15.56) | 0.118 | 0.549 | 0.710 |
|  | FP | 17.84  (2.37) | 18.25  (2.25) | 3.10  (11.98) | 0.338 | 0.181 | 0.404 |
| Plasma | Free-fraction | 0.24  (0.02) | 0.25  (0.02) | 6.36  (12.77) | 0.139 | 0.450 | 0.588 |

*Supplementary Table 4. Changes in [^11^C]-UCBJ metrics before and 7-8 days after ketamine, Day 7-8 group (N = 6). The table report the average VT, DVR-1, and FP values within the pre-defined ROIs. The p-values are the result of one-sided linear mixed-effect model with α significance threshold set at 0.05. The effect size was computed via Cohen’s d test. The statistical confidence of the result was computed via BF analysis.*

7.4. Correlation between [^11^C]-UCBJ metrics

At baseline, the VT and FP [^11^C]-UCBJ PET metrics were positively correlated in all analysed ROIs. No correlation was found between [^11^C]-UCBJ DVR-1 and the other PET metrics. After ketamine, there were no correlations between the PET metrics. All results are shown in Supplementary Table 5. There was no correlation across changes in [11C]-UCBJ VT, DVR-1, and FP (Supplementary Table 6).

| All participants (N = 11) | | | | | | |
| --- | --- | --- | --- | --- | --- | --- |
| ROIs | Timepoint | [^11^]-UCBJ Metric 1 | [^11^]-UCBJ Metric 2 | R coefficient | p value | p adj. |
| dlPFC | Scan 1 | VT | DVR-1 | 0.48 | 0.137 | 0.231 |
|  | Scan 1 | VT | FP | 0.9 | < 0.001 | 0.001 |
|  | Scan 1 | FP | DVR-1 | 0.29 | 0.386 | 0.386 |
|  | Scan 2 | VT | DVR-1 | 0.44 | 0.183 | 0.235 |
|  | Scan 2 | VT | FP | 0.45 | 0.173 | 0.235 |
|  | Scan 2 | FP | DVR-1 | 0.35 | 0.299 | 0.317 |
| vmPFC | Scan 1 | VT | DVR-1 | 0.46 | 0.154 | 0.231 |
|  | Scan 1 | VT | FP | 0.94 | <0.001 | <0.001 |
|  | Scan 1 | FP | DVR-1 | 0.43 | 0.193 | 0.240 |
|  | Scan 2 | VT | DVR-1 | 0.45 | 0.163 | 0.235 |
|  | Scan 2 | VT | FP | 0.5 | 0.121 | 0.235 |
|  | Scan 2 | FP | DVR-1 | 0.35 | 0.286 | 0.317 |
| ACC | Scan 1 | VT | DVR-1 | 0.46 | 0.154 | 0.232 |
|  | Scan 1 | VT | FP | 0.87 | <0.001 | 0.003 |
|  | Scan 1 | FP | DVR-1 | 0.45 | 0.173 | 0.239 |
|  | Scan 2 | VT | DVR-1 | 0.45 | 0.173 | 0.235 |
|  | Scan 2 | VT | FP | 0.51 | 0.114 | 0.235 |
|  | Scan 2 | FP | DVR-1 | 0.47 | 0.146 | 0.235 |
| PCC | Scan 1 | VT | DVR-1 | 0.41 | 0.214 | 0.241 |
|  | Scan 1 | VT | FP | 0.89 | <0.001 | 0.002 |
|  | Scan 1 | FP | DVR-1 | 0.34 | 0.313 | 0.331 |
|  | Scan 2 | VT | DVR-1 | 0.55 | 0.082 | 0.235 |
|  | Scan 2 | VT | FP | 0.59 | 0.061 | 0.235 |
|  | Scan 2 | FP | DVR-1 | 0.58 | 0.066 | 0.235 |
| Amygdala | Scan 1 | VT | DVR-1 | 0.57 | 0.071 | 0.182 |
|  | Scan 1 | VT | FP | 0.86 | 0.001 | 0.004 |
|  | Scan 1 | FP | DVR-1 | 0.47 | 0.146 | 0.232 |
|  | Scan 2 | VT | DVR-1 | 0.45 | 0.173 | 0.235 |
|  | Scan 2 | VT | FP | 0.45 | 0.163 | 0.235 |
|  | Scan 2 | FP | DVR-1 | 0.44 | 0.183 | 0.235 |
| Hippocampus | Scan 1 | VT | DVR-1 | 0.5 | 0.121 | 0.232 |
|  | Scan 1 | VT | FP | 0.93 | < 0.001 | < 0.001 |
|  | Scan 1 | FP | DVR-1 | 0.41 | 0.214 | 0.232 |
|  | Scan 2 | VT | DVR-1 | 0.41 | 0.214 | 0.257 |
|  | Scan 2 | VT | FP | 0.55 | 0.087 | 0.235 |
|  | Scan 2 | FP | DVR-1 | 0.16 | 0.634 | 0.634 |

*Supplementary Table 5:* *Correlations between [^11^C]-UCBJ metrics at Scan 1 and at Scan 2, all participants (N = 11). The correlation coefficients (R) and p-values results from pair-wise spearman correlations between the PET metrics. The p adj. are the results of FDR correction using the Benjamini-Hochberg adjustment.*

| ROIs | Δ [^11^]-UCBJ Metric 1 | Δ [^11^]-UCBJ Metric 2 | R coefficient | p value | p adj. |
| --- | --- | --- | --- | --- | --- |
| dlPFC | VT | DVR-1 | 0.6 | 0.056 | 0.275 |
|  | VT | FP | 0.41 | 0.214 | 0.331 |
|  | FP | DVR-1 | 0.47 | 0.146 | 0.275 |
| vmPFC | VT | DVR-1 | 0.63 | 0.044 | 0.275 |
|  | VT | FP | 0.45 | 0.163 | 0.278 |
|  | FP | DVR-1 | 0.36 | 0.273 | 0.333 |
| ACC | VT | DVR-1 | 0.5 | 0.121 | 0.275 |
|  | VT | FP | 0.42 | 0.203 | 0.321 |
|  | FP | DVR-1 | 0.35 | 0.299 | 0.333 |
| PCC | VT | DVR-1 | 0.60 | 0.056 | 0.275 |
|  | VT | FP | 0.48 | 0.137 | 0.275 |
|  | FP | DVR-1 | 0.39 | 0.237 | 0.336 |
| Amygdala | VT | DVR-1 | 0.54 | 0.094 | 0.275 |
|  | VT | FP | 0.49 | 0.129 | 0.275 |
|  | FP | DVR-1 | 0.35 | 0.299 | 0.333 |
| Hippocampus | VT | DVR-1 | 0.35 | 0.299 | 0.275 |
|  | VT | FP | 0.31 | 0.356 | 0.356 |
|  | FP | DVR-1 | 0.34 | 0.313 | 0.333 |

*Supplementary Table 6:* *Correlations between change in [^11^C]-UCBJ metrics from Scan 1 to Scan 2, all participants (N = 11). The correlation coefficients (R) and p-values results from pair-wise spearman correlations between the PET metrics. The p adj. are the results of FDR correction using the Benjamini-Hochberg adjustment.*

7.5. Correlation between [^11^C]-UCBJ and Sleep

Upon arrival at the scanning facility, participants reported their estimated sleep duration for the night preceding each PET/MRI scan (before and after ketamine). No significant correlations were observed between [^11^C]-UCBJ and the hours of sleep preceding the relative PET/MRI scan (Data not shown). A positive relationship was observed between Δ [^11^C]-UCBJ VT and Δ Sleep hours in all ROIs, reaching significance in the vmPFC and the PCC. This would suggest that participants who slept more before the post-ketamine scan also showed relative increases in [^11^C]-UCBJ VT in the vmPFC and PCC. However, these correlations they did not survive FDR correction. These data suggest a potential effect of sleep on [^11^C]-UCBJ VT which warrants for further exploration through a quantitative assessment of sleep patterns.

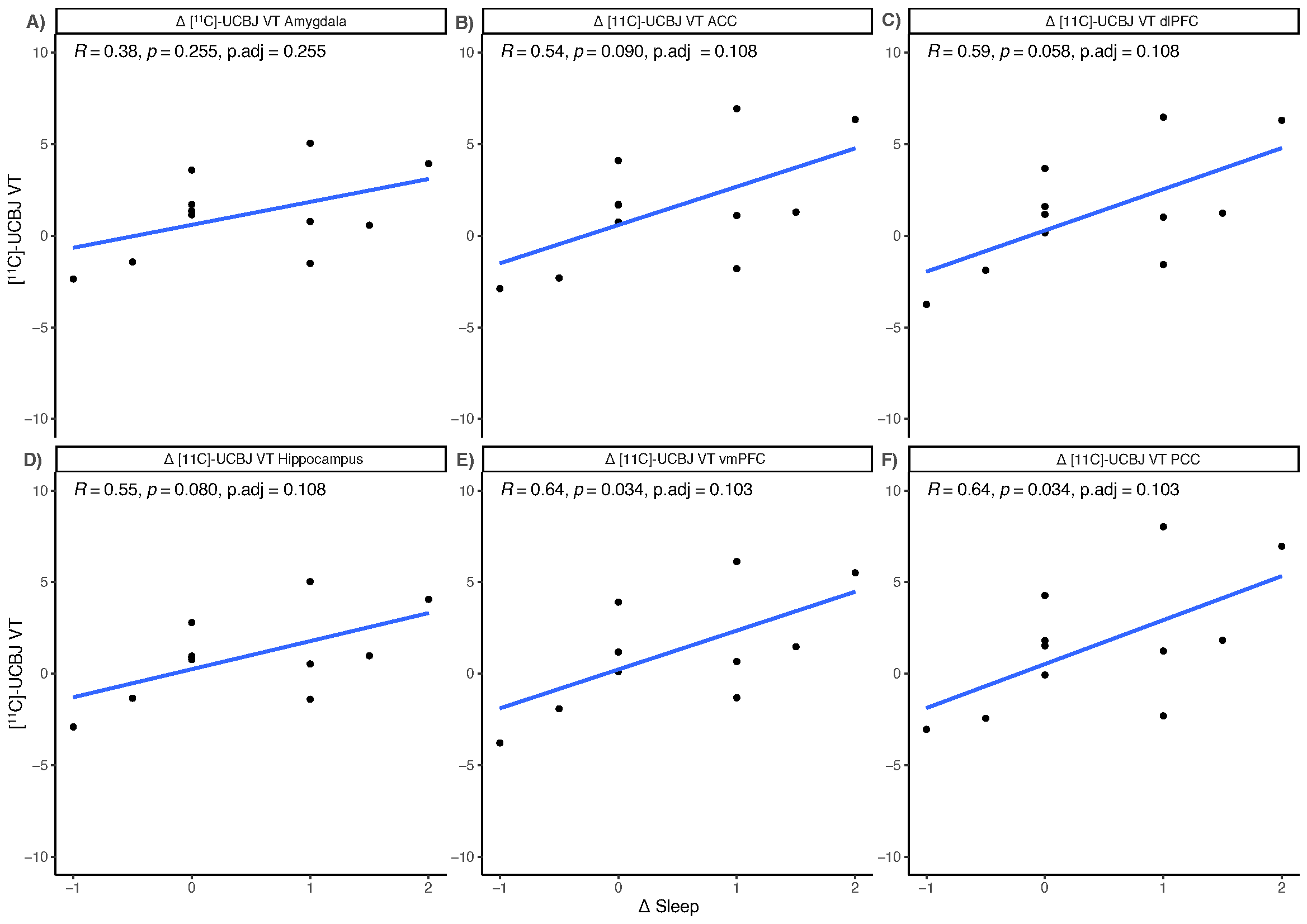
*Supplementary Figure 1. Correlations between Δ [^11^C]-UCBJ VT and Δ sleep. The plot shows Spearman’s correlations between the difference in [^11^C]-UCBJ VT between the 2 scans and the difference in amount of sleep before the 2 scans in the pre-defined ROIs (A-F). The p adj. are the results of FDR correction using the Benjamini-Hochberg adjustment.*

8. Ketamine effects on ^1^H-MRS metrics

Supplementary Table 7 shows the change in ^1^H-MRS-derived metrics following ketamine administration in the ACC.

| ACC PRESS sequence (n = 9) | | | | | | | |
| --- | --- | --- | --- | --- | --- | --- | --- |
| Correction | MRS Metric | Scan 1 (SD) | Scan 2 (SD) | % Difference (SD) | p-value | Cohen’s d | Bayes factor |
| Tissue-corrected | Glutamate | 14.60  (2.48) | 18.68  (3.35) | 21.77  (25.35) | 0.030 | 0.878 | 2.70 |
|  | Glutamine | 7.16  (3.65) | 6.10  (2.64) | 93.97  (295.80) | 0.493 | -0.229 | 0.391 |
|  | Glutamine/Glutamine index | 21.70  (4.85) | 23.53  (5.60) | 13.52  (36.62) | 0.456 | 0.261 | 0.414 |
| Creatine- corrected | Glutamate | 1.21  (0.09) | 1.21  (0.11) | -0.48  (7.03) | 0.815 | -0.081 | 0.330 |
|  | Glutamine | 0.62  (0.36) | 0.42  (0.13) | 77.22  (310.20) | 0.140 | -0.547 | 0.865 |
|  | Glutamine/Glutamine index | 1.83  (0.41) | 1.62  (0.19) | -8.12  (19.01) | 0.144 | -0.540 | 0.847 |
| ACC HERMES sequence (n = 6) | | | | | | | |
| Correction | MRS Metric | Scan 1 (SD) | Scan 2 (SD) | % Difference (SD) | p-value | Cohen’s d | Bayes factor |
| Tissue-corrected | GABA | 0.55  (0.35) | 0.40  (0.11) | -11.22  (38.06) | 0.342 | -0.401 | 0.540 |
|  | GSH | 0.29  (0.19) | 0.49  (0.30) | 1488.33  (3599.75) | 0.170 | 0.654 | 0.881 |
| Creatine- corrected | GABA | 0.31  (0.17) | 0.40  (0.11) | 56.42  (79.19) | 0.289 | 0.433 | 0.570 |
|  | GSH | 0.15  (0.10) | 0.26  (0.18) | 1391.60  (3373.70) | 0.220 | 0.573 | 0.745 |

*Supplementary Table 7. Changes in ^1^H-MRS metrics before and after ketamine. The table report the average values extracted from the PRESS and HERMES sequences within the ACC. The p-values are the result of two-sided linear mixed-effect model with α significance threshold set at 0.05. The effect size was computed via Cohen’s d test. The statistical confidence of the result was computed via BF analysis.*

9. Ketamine effects on miFC

Ketamine effects on within-network miFC are shown in Supplementary Table 7.

| **All subjects miFC (n = 11)** | | | | |
| --- | --- | --- | --- | --- |
| **PET/MRI-2 > PET/MRI-1** | |  | **PET/MRI-1 > PET/MRI-2** | |
| Proportion | Degree | Network | Degree | Proportion |
| 0.14 | 507 | Visual | 823 | 0.12 |
| 0.07 | 251 | Somatomotor | 1065 | 0.16 |
| 0.12 | 450 | Dorsal Attention | 660 | 0.1 |
| 0.12 | 452 | Salience/Ventral Attention | 708 | 0.11 |
| 0.07 | 227 | Limbic | 455 | 0.07 |
| 0.22 | 807 | Control | 1431 | 0.21 |
| 0.24 | 873 | DMN | 1417 | 0.21 |
| 0.03 | 113 | Temporal Parietal | 181 | 0.03 |

*Supplementary Table 8. The network-level effects of ketamine on fMRI-derived miFC. Degree is a graph-theoretic term which refers to the total number of regions within a network showing significantly different miFC between sessions. The proportion refers to the degree of that network divided by the sum of degree for all networks, i.e., the percent of significant changes in regional miFC contained within that network.*
